# Yap suppresses T cell function and infiltration in the tumor microenvironment

**DOI:** 10.1101/644757

**Authors:** Eleni Stampouloglou, Anthony Federico, Emily Slaby, Stefano Monti, Gregory L. Szeto, Xaralabos Varelas

## Abstract

A major challenge for cancer immunotherapy is sustaining T cell activation and recruitment in immunosuppressive solid tumors. Here we report that Yap levels are sharply induced upon activation of CD4^+^ and CD8^+^ T cells and that Yap functions as an immunosuppressive factor and inhibitor of effector differentiation. Loss of Yap in T cells results in enhanced T cell activation, differentiation and function, which translates *in vivo* to an improved ability for T cells to infiltrate and repress tumors. Gene expression analyses of tumor-infiltrating T cells following Yap deletion implicates Yap as a mediator of global T cell responses in the tumor microenvironment and as a key negative regulator of T cell tumor infiltration and patient survival in diverse human cancers. Collectively, our results indicate that Yap plays critical roles in T cell biology, and suggest that inhibiting Yap activity improves T cell responses in cancer.

## INTRODUCTION

CD8^+^ and CD4^+^ T cells are central players in the adaptive immune system. T cells elicit targeted, antigen-specific responses for direct killing of an infected or transformed cell, shaping and regulating the immune response in host defense [1]. Most mature T cells circulate in a resting, naïve state, and upon cognate antigen recognition, T cells become activated, proliferate clonally, and differentiate into effector T cells. Naïve CD8^+^ T cells differentiate into cytotoxic T cells, while CD4^+^ T cells differentiate into an array of different types of helper T cells (i.e., Th1, Th2, Th17 or Treg) depending on microenvironmental cues [2]. Each phenotype is defined by expression of signature transcription factors and effector cytokines leading to distinct functions [3].

T cell activation also upregulates negative feedback mechanisms, such as inhibitory receptors, which restrain their action to minimize pathogenic inflammation and autoimmunity [1, 4]. This network of immunosuppressive factors is frequently co-opted in chronic infections and cancer, leading to terminally differentiated and exhausted T cells that lose effector function and ability to infiltrate disease sites [5]. The finding that revitalization of exhausted, dysfunctional T cells can restore the immune response has revolutionized cancer therapy with the use of checkpoint inhibition [6]. Chimeric antigen receptor T cells (CAR-T), engineered for enhanced antigen recognition and co-stimulation also demonstrate promising clinical efficacy [7–9]. However, both immunotherapies are effective for only a fraction of patients [10–14]. Major challenges to extending the efficacy of immunotherapy to more cancer patients include sustaining T cell activation and achieving T cell infiltration in the immunosuppressive microenvironment of solid tumors [15–24]. Improved understanding of mechanisms controlling T cell differentiation and function is critical to overcoming these barriers.

Yap is a key effector of the Hippo signaling pathway, directing transcriptional programs that control stem cell biology by integrating microenvironmental and cell intrinsic cues [25]. Yap regulation leads to control of cell death and survival, proliferation and cell fate determination, and dysregulated Yap activity contributes to disease, most notably cancer [26]. While the dynamics of Yap regulation coupled with differentiation are well characterized in stem cells and tissue specific progenitor cells, less is known about the role of Yap in T cells. The Hippo pathway has been implicated in coupling CD8^+^ T cell clonal expansion to terminal differentiation through upregulation of kinase LATS1 and Yap degradation [27]. The Hippo pathway kinases MST1/2 have also been implicated in thymocyte egress and antigen recognition, lymphocyte polarization, adhesion and trafficking, survival, differentiation and proliferation [28–36]. Further, an immune-cell intrinsic role for Yap in CD4^+^ T cells has also been described, with Yap being shown to be critical for potentiating TGFβ/SMAD signaling responses that direct Treg function [37]. In this study, the conditional deletion of the Yap gene using CD4-Cre or Foxp3-Cre models was shown to lead to reduced tumor growth in syngeneic mouse models of cancer, with Yap being linked mechanistically to Treg function and stability *in vitro* and *in vivo* [37]. However, the involvement of Yap in CD8^+^ T cells and other CD4^+^ subtypes has remained understudied.

In this study, we sought to determine whether Yap has overlapping control over both CD4^+^ and CD8^+^ T cells and how these functions may affect cancer immunity. We observed that Yap expression is elevated upon T cell activation, and that conditional deletion of the *Yap* gene in CD4^+^ and CD8^+^ T cells using the CD4-Cre model leads to enhanced T-cell activation and differentiation. These phenotypes translated to the reduced growth of B16F10 and LLC tumors in CD4Cre Yap-deleted mice, which showed strongly increased T cell tumor infiltration. Interestingly, using adoptive cell transfer experiments we observed that Yap-deleted CD8^+^ T cells have an intrinsic ability to infiltrate tumors with much higher efficiency, which is a major goal for improving cancer immunotherapy. RNA-sequencing analyses of Yap-deleted tumor infiltrated lymphocytes (TILs) revealed upregulation of key signaling pathways in T cell activation, differentiation and function in CD4^+^ and CD8^+^ T cells. Notably, we found that Yap regulated gene expression changes were tumor-specific, as we observed minimal gene expression changes in lymphocytes isolated tumor-draining lymph nodes. Yap-regulated gene expression signatures from TILs correlated with T cell infiltration and patient survival across multiple human cancers in The Cancer Genome Atlas (TCGA), including melanoma and lung cancer, as demonstrated in our mouse studies. Our study merges new evidence with observations from prior studies, highlighting Yap as a broad suppressor of CD4^+^ and CD8^+^ T cell activation and function and a key regulator of T cell tumor infiltration and survival in cancer immunotherapy patients.

## RESULTS

### Yap inhibits CD4^+^ and CD8^+^ T cell activation

To study the role of Yap in T cells, we started by analyzing Yap levels in isolated primary CD4^+^ and CD8^+^ T cells from mice that were either unstimulated or activated with anti-CD3/CD28 coated beads for 24 hours. We observed that *Yap* RNA levels were dramatically increased upon stimulation of CD4^+^ and CD8^+^ T cells (Figure 1A-B). This observation contrasted prior reports that Yap expression is exclusive to activated Tregs or CD8^+^ T cells cultured under specific conditions [27, 37], which prompted us to explore more broad roles for Yap in T cell activation and function. To gain insight into the roles of Yap in T cells, we generated a mouse model whereby Yap deletion and EYFP expression are induced under the control of the CD4 promoter (Yap-loxP/loxP: LSL-EYFP; CD4-Cre, herein referred to as Yap-cKO). No systemic defects or gross phenotypes were observed in Yap-cKO mice housed in a barrier facility. Deletion of the *Yap* gene was achieved in both CD4^+^ and CD8^+^ T cells at the CD4^+^CD8^+^ double-positive stage of T cell development, as confirmed by EYFP expression (Supplementary Figure 1A-B).

**Figure 1.**
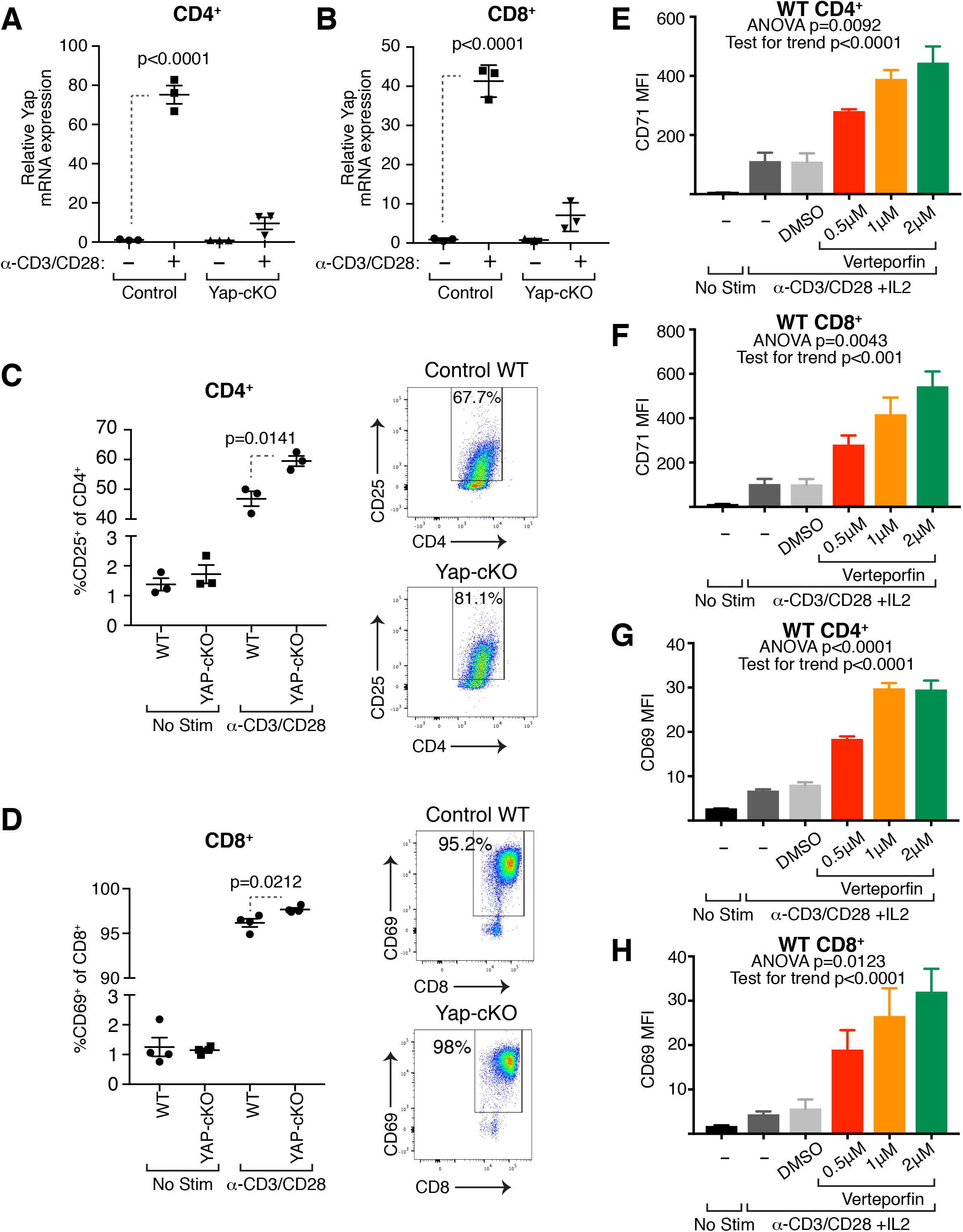
Yap RNA expression is induced upon T cell activation resulting in suppression of T cell activation. Wild type (WT) and YAP-cKO CD4^+^ and CD8^+^ T cells were isolated from mouse spleens and stimulated with *α*CD3 and *α*CD28 coated magnetic beads to test expression of YAP mRNA and activation markers. Activation marker expression was also tested in WT CD4^+^ and CD8^+^ T cells treated with increasing concentrations of Verteporfin under IL-2, *α*CD3 and *α*CD28 stimulation. Statistical differences were determined by using a Student’s t-test (a-d) or one-way repeated-measures ANOVA with post-hoc test for linear trend with increasing verteporfin (e-h). a. YAP mRNA expression by CD4^+^ T cells isolated from WT or YAP-cKO mice 24 hours post CD3/CD28 stimulation (n=3/group). b. YAP mRNA expression by CD8^+^ T cells isolated from WT or YAP-cKO mice 24 hours post CD3/CD28 stimulation (n=3/group). c. CD25 expression on WT and YAP-cKO CD4^+^ T cells 24 hours post CD3/CD28 stimulation (n=3/group). d. CD69 expression on WT and YAP-cKO CD8^+^ T cells 24 hours post CD3/CD28 stimulation (n=3/group). e. CD71 expression on WT CD4^+^ T cells three days post IL-2 and CD3/CD28 stimulation and increasing concentration of verteporfin (n=4/group). f. CD71 expression on WT CD8^+^ T cells three days post IL-2 and CD3/CD28 stimulation and increasing concentration of verteporfin (n=4/group). g. CD69 expression on WT CD4^+^ T cells three days post IL-2 and CD3/CD28 stimulation and increasing concentration of verteporfin (n=4/group). h. CD69 expression on WT CD8^+^ T cells three days post IL-2 and CD3/CD28 stimulation and increasing concentration of verteporfin (n=4/group).

We next measured the surface levels of activation markers (CD69 and CD25) in Yap-cKO T cells. CD4^+^ and CD8^+^ T cells were isolated from wild type (WT) and Yap-cKO mouse spleens, activated, and levels of CD69 and CD25 were measured 24 hours later. We found increased levels of CD25 in Yap-cKO CD4^+^ T cells (Figure 1C-D) and CD69 in Yap-cKO CD8^+^ T cells (Figure 1E-F) compared to WT cells. CD4^+^ and CD8^+^ T cell proliferation was also measured, but no significant differences were observed between WT and Yap-cKO cells 72 hours post-stimulation (Supplementary Figure 1E-F). We confirmed the Yap-cKO effects in wild-type T cells by using the small molecule drug verteporfin (also known as Visudine) which is a known inhibitor of Yap binding to TEADs [38]. Verteporfin treatment of CD4^+^ and CD8^+^ T cells isolated from WT mice increased levels of early activation markers CD71 (Figure 1E-F) and CD69 (Figure 1G-H) in a concentration-dependent manner. These increases were sustained up to 3 days post-activation, suggesting verteporfin or Yap deletion can enhance T cell activation. Consistent with the phenotype observed with Yap-cKO cells, verteporfin treatment did not significantly affect proliferation of T cells isolated from WT mice 3 days post stimulation (Supplementary Figure 1G). Together, these data indicate that Yap is induced upon T cell activation and functions as a suppressor of T cell activation.

### Yap inhibits CD4^+^ T cell differentiation

Upon encountering their cognate antigen and receiving appropriate co-stimulation, naïve CD4^+^ T cells become activated and differentiate into several subsets, including Th1, Th2 and Th17, to contribute to protective immunity or immunopathology depending on microenvironmental signals [3]. Signature transcription factors and effector cytokines define each subset: Th1 is defined by expression of T-BET and IFN*γ*, Th2 by GATA3 and IL-4, and Th17 by ROR*γ*t and IL-17. Given the wealth of evidence for Yap in stem cell regulation and the coordination of Yap inactivation with cell differentiation, we hypothesized that deletion of Yap alters the ability of CD4^+^ T cells to differentiate. To test this hypothesis, we isolated naïve CD4^+^ T cells from WT and Yap-cKO mice and cultured them under Th1, Th17 and Th2 polarizing conditions [39, 40]. We found that Yap deleted CD4^+^ T cells showed significantly enhanced Th1, Th17, and Th2 differentiation compared to WT cells, demonstrated by increased intracellular IFN*γ*, IL-17, and GATA3 expression, respectively (Figure 2A-C). These observations were distinct from prior findings that suggested Treg-specific functions for Yap [37], and implicate Yap as a broad inhibitor of CD4^+^ T cell activation and differentiation.

**Figure 2.**
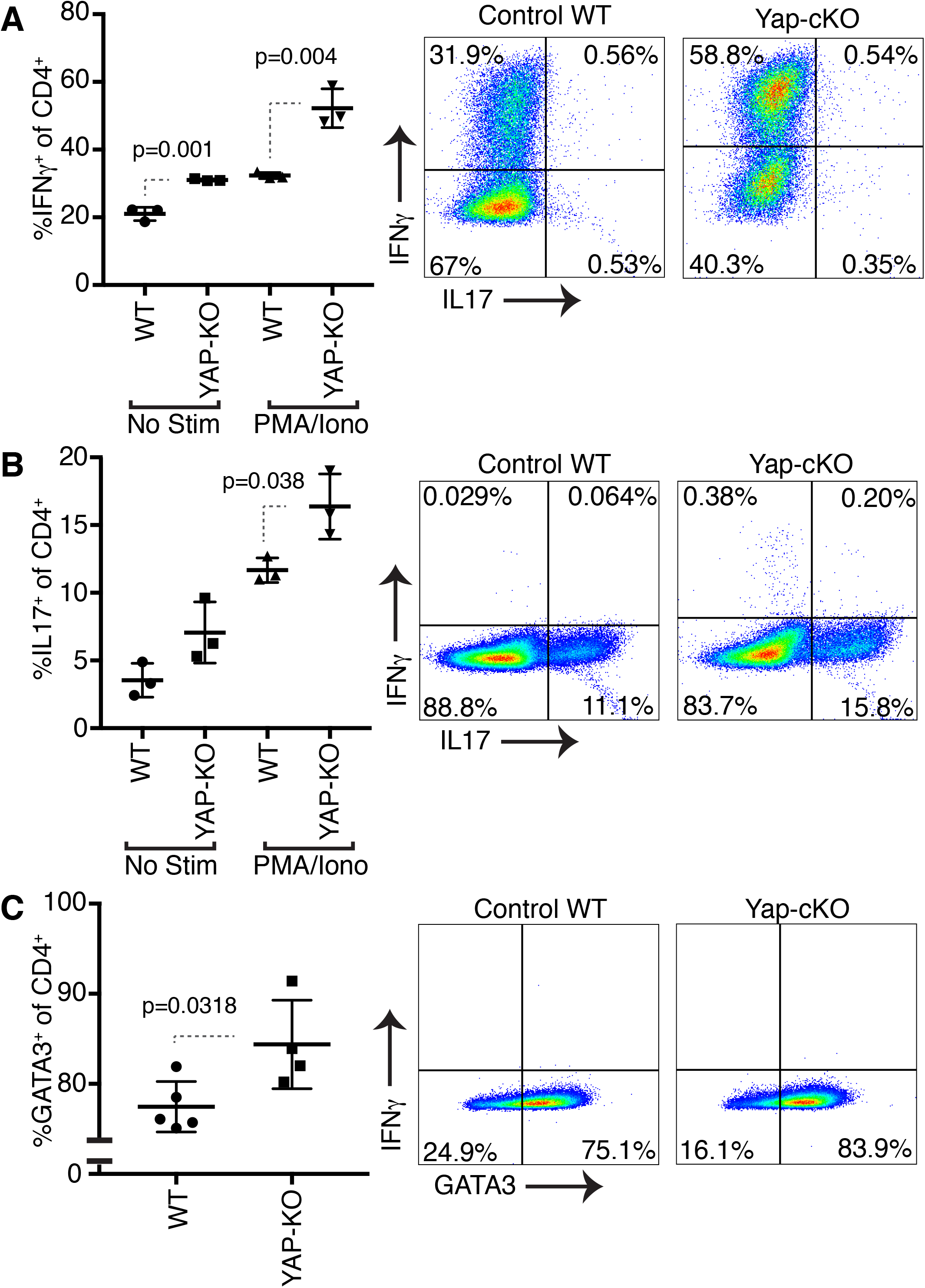
Deletion of YAP in CD4^+^ T cells results in increased IFNγ, IL-17 and GATA3 expression under Th1, Th17 and Th2 polarizing conditions, respectively. Naïve CD4^+^ T cells from WT and YAP-cKO mice were isolated using magnetic beads. WT and YAP KO T cells were cultured under Th1, Th17 or Th2 skewing conditions for 5 days in the presence of CD3 and CD28 antibodies. On Day5, IFN*γ*, IL-17 and GATA3 expression was measured using flow cytometry. Statistical differences were determined by using a Student’s t-test. a. IFN*γ* and IL-17 expression in WT and YAP-cKO CD4^+^ T cells under Th1 polarizing conditions (n=3/group). b. IFN*γ* and IL-17 expression in WT and YAP-cKO CD4^+^ T cells under Th17 polarizing conditions (n=3/group). c. IFN*γ* and GATA3 expression in WT and YAP-cKO CD4^+^ T cells under Th2 polarizing conditions (n=3/group).

### Deletion of Yap in T cells promotes T cell infiltration into developing tumors and blocks tumor growth

Since we observed that Yap inhibits T cell activation and differentiation in vitro, we aimed to study the effect of Yap deletion in vivo. We started by examining the growth of the subcutaneous B16F10 mouse tumor model in Yap-cKO mice, due to the poor immunogenic phenotype and highly immunosuppressive microenvironment that leads to low T cell infiltration in this model [41–44]. Yap deletion in T cells resulted in superior anti-tumor immunity, as evidenced by the significant delay in tumor growth in Yap-cKO compared to WT mice (Figure 3A-B), consistent with prior observations [37]. The growth of subcutaneous Lewis lung carcinoma (LLC) tumors was also significantly reduced in Yap-cKO mice (Figure 3C-D), suggesting a general role for Yap in anti-tumor immune responses.

**Figure 3.**
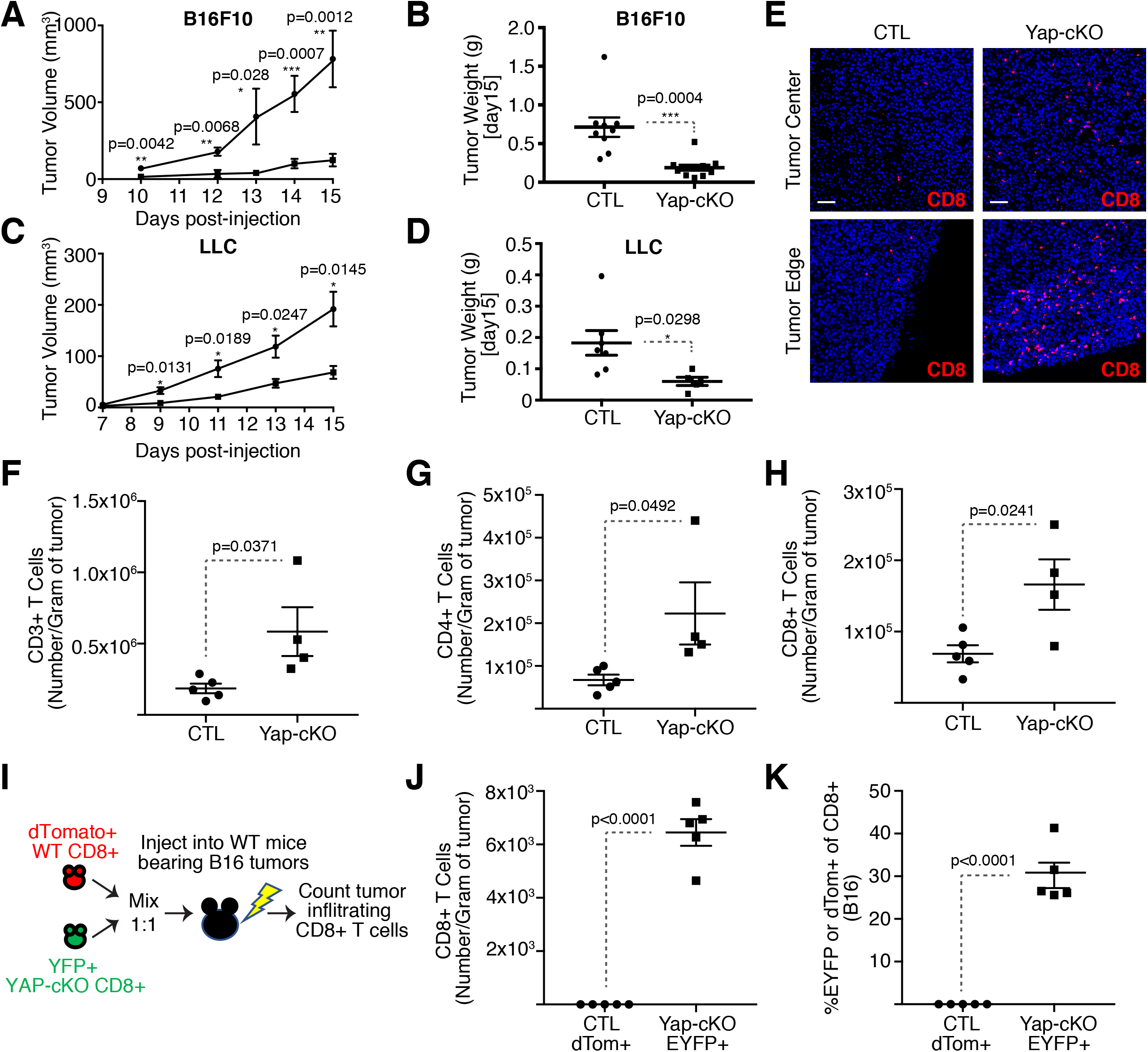
T cell specific deletion of YAP results in reduced tumor growth and enhanced T cell tumor infiltration. Mice were challenged subcutaneously with B16F10 or LLC cells on the right flank. Some mice carrying B16F10 tumors received adoptive cell transfer of WT and YAP-cKO CD8^+^ T cells. Tumor growth was monitored over the course of 15 days, until the maximum size of the tumors reached 500mm3. B16F10 tumors were harvested for immunofluorescence or flow cytometric analysis. Statistical differences were determined by using a Student’s t-test. a. B16 tumor growth curve of WT and YAP-cKO mice (n=9/group). b. Tumor weight of B16 tumors derived from WT and YAP-cKO mice on Day 15 post injection (n=9/group). c. LLC tumor growth curve of WT and YAP-cKO mice (n=7/WT group, n=5/cKO group). d. Tumor weight of LLC tumors derived from WT and YAP-cKO mice on Day 15 post injection (n=7/WT group, n=5/cKO group). e. CD8+ T cell immunofluorescence on day 15 of B16 tumor growth. f. Absolute numbers of CD3^+^ TILs from WT and YAP-cKO B16 tumors. Tumors were harvested on Day 15, stained with antibodies against CD45, CD3, CD4 and CD8 and analyzed using flow cytometry (n=5/WT mice, n=4/ YAP-cKO mice). g. Absolute numbers of CD4^+^ TILs from WT and YAP-cKO B16 tumors, prepared as in Fig4d (n=5/WT mice, n=4/ YAP-cKO mice). h. Absolute numbers of CD8+ TILs from WT and YAP-cKO B16 tumors, prepared as in Fig4d (n=5/WT mice, n=4/ YAP-cKO mice). i. Experimental plan for adoptive cell transfer of WT and YAP-cKO CD8+ T cells in WT B16 bearing mice. j. Absolute numbers of dTom+ and EYFP+ WT vs YAP-cKO CD8+ T cells in C57BL/6 B16 tumors. WT dTom+ CD8+ T cells were mixed 1:1 with YAP-cKO EYFP+ CD8+ T cells prior to being injected into WT C57BL/6 mice. Subsequently, mice were injected subcutaneously with B16F10 melanoma cells and absolute number of infiltrating T cells was determined on D15 by flow cytometry (n=5/group). k. Percentage of dTom^+^ and EYFP^+^ CD8^+^ T cells out of total B16 tumor infiltrating CD8^+^ T cells (n=5/group).

Given the strong correlation between CD8^+^ T cell tumor infiltration and patient survival as well as response to immunotherapy [15-19, 45-49], we investigated the extent of T cell infiltration in tumors developed in WT versus Yap-cKO mice. Immunofluorescence microscopy analysis revealed that tumors from Yap-cKO mice were significantly more infiltrated with CD8^+^ T cells at both the tumor center and tumor edge compared to WT mice (Figure 3E). Flow cytometry revealed more Yap-cKO CD4^+^ and CD8^+^ T cells infiltrating tumors compared to WT counterparts (Figure 3F-H).

To directly address whether Yap-deleted T cells have an increased ability for tumor infiltration we isolated CD8^+^ T cells from Yap-cKO and WT mice and directly compared them in adoptive transfer experiments in WT mice challenged with B16F10 tumor cells (illustrated in Figure 3I). For this, we isolated equal numbers of tdTomato^+^ CD8^+^ T cells from WT mice and EYFP^+^ CD8^+^ T cells from Yap-cKO mice and mixed these cells at a 1 to 1 ratio. Cell mixtures were intravenously injected into WT mice the same day as subcutaneous injection of B16F10 cells. Absolute numbers of tdTomato^+^ (WT) and EYFP^+^ (Yap-cKO) CD8^+^ T cells were then measured in tumors after 15 days. Yap-cKO CD8^+^ T cells were significantly infiltrated in tumors (Figure 3J), with nearly 30% of all CD8^+^ tumor-infiltrating T cells being EYFP^+^ Yap-cKO T cells compared to almost no WT tdTomato^+^ T cells from being observed in tumors (Figure 3K). These results conclusively show that Yap-cKO CD8^+^ T cells have enhanced intrinsic tumor infiltration capacity.

### Yap regulates global T cell responses in the local tumor microenvironment

We next aimed to define Yap-regulated signaling networks in T cells that impacted tumor growth and T cell infiltration. CD4^+^ and CD8^+^ TILs were isolated from B16 tumors grown in WT or Yap-cKO mice, as well as CD4^+^ and CD8^+^ T cells from corresponding tumor draining lymph nodes (TDLNs), and analyzed by RNA-Seq **(Supplementary Tables 1-3)**. A large number of genes were differentially expressed in CD4^+^ and CD8^+^ TILs isolated from Yap-cKO mice compared to WT mice (Figure 4A-B and Supplementary Figure 2A-B). Notably, T cells in the TDLNs showed markedly fewer gene expression changes with Yap deletion than those observed with TILs (Figure 4C-D). These data are consistent with Yap function being coordinated with T cell activation, and suggest that the enhanced anti-tumor responses observed in Yap-cKO T cells are mediated by changes in cellular responses to local tumor microenvironment signals.

**Figure 4.**
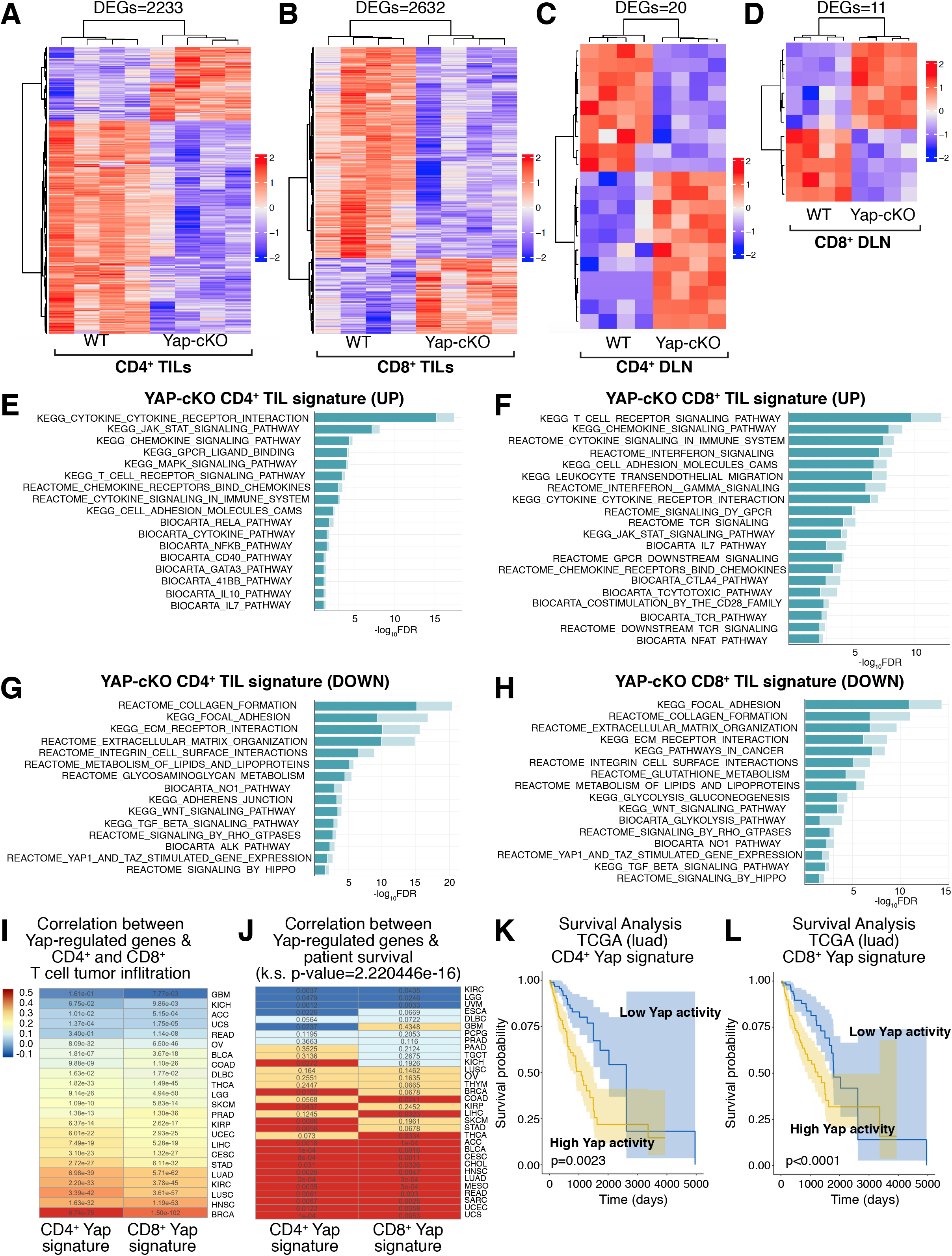
RNA-Seq analysis of WT and CD4Cre YAP KO CD4^+^ and CD8^+^ B16 TILs uncovers YAP as a negative regulator of T cell activation, differentiation, cytokine signaling and migration. a. WT vs YAP-cKO CD4^+^ TIL DEG heatmap. b. WT vs YAP-cKO CD8^+^ TIL DEG heatmap. c. WT vs YAP-cKO CD4^+^ TDLN DEG heatmap. d. WT vs YAP-cKO CD8^+^ TDLN DEG heatmap. e. Hyper-enrichment analysis shows enrichment of induced gene sets observed in CD4^+^ YAP-cKO TILs. f. Hyper-enrichment analysis shows enrichment of induced gene sets observed in CD8^+^ YAP-cKO TILs. g. Hyper-enrichment analysis shows enrichment of repressed gene sets observed in CD4^+^ YAP-cKO TILs. h. Hyper-enrichment analysis shows enrichment of repressed gene sets observed in CD8^+^ YAP-cKO TILs. i. Gene expression changes identified in YAP-cKO CD4^+^ and CD8^+^ TILs correlate with genes reflecting tumor infiltration across many cancers in TCGA data. The heatmap is colored by the coefficient and the text of each cell represents the adjusted p-value of the correlation. j. Gene expression changes identified in YAP-cKO CD4^+^ and CD8^+^ T cells correlate with patient survival data available in TCGA across several cancers. Blue: the average survival probability for patients with low activity of the signature is higher. Red: the average survival probability for patients with high activity of the signature is higher. Yellow: neither group has a higher probability of survival. The text of each cell is the p-value for the survival estimation. The distribution of p-values arising from the multiple survival analyses for each signature across TCGA datasets was compared to a uniform distribution using a Kolmogorov-Smirnov test. k. Kaplan-Meier Survival analysis showing the average survival probability of patients with lung adenocarcinoma that show low versus high YAP activity derived from the YAP-cKO CD4^+^ gene expression signature. l. Kaplan-Meier Survival analysis showing the average survival probability of patients with lung adenocarcinoma that show low versus high YAP activity derived from the YAP-cKO CD8^+^ gene expression signature.

Hyper-enrichment analyses [50] of the gene expression changes identified in TILs from Yap-cKO mice showed an induction of genes related to T cell activation, differentiation, survival and migration (Figure 4E-F) (**Supplementary Table 3**). Yap-cKO TILs show enrichment of genes associated with T cell receptor signaling, genes that encode major co-stimulatory molecules, and genes downstream of TCR signaling (Supplementary Figure 3A-B), suggesting enhanced responsiveness to TCR signals. Cytokine and cytokine receptor signaling gene sets are also enriched in Yap-cKO TILs, suggesting that cytokine production as well as responsiveness to cytokine receptor engagement are enhanced. Further, chemokine and chemokine receptor signaling gene sets are enriched in Yap-cKO TILs (Supplementary Figure 3C-F), which likely contributes to their improved tumor infiltrating capacities. Subset defining transcription factors and cytokines associated with each of the major CD4^+^ T cell phenotypes were all upregulated in Yap KO CD4^+^ TILs (Supplementary Figure 3G-H), suggesting that Yap deletion leads to enhanced naive CD4^+^ T cell differentiation. Using unique upregulated genes after differentiation to each of the major T helper subsets [51], we show that Yap-cKO CD4^+^ TILs are more skewed towards a Th2 and Treg phenotype compared to WT CD4^+^ TILs (Supplementary Figure 4A-D). As expected, gene set enrichment profiles identified in Yap-cKO TILs also included “Yap1 and TAZ stimulated gene expression” and “signaling by Hippo”, which were repressed in both CD4^+^ and CD8^+^ TILs (Figure 4G-H). Altogether, these analyses demonstrate that Yap-cKO T cells may have potentiated signaling from TCR and cytokines due to increased expression of key signaling adaptors and pathway molecules. These insights suggest that Yap function, even in Tregs, may regulate more fundamental signaling processes that are distinct from those suggested in prior studies [37].

Comparison of gene expression changes identified in CD4^+^ and CD8^+^ Yap-cKO TILs with clinical data from TCGA [52] showed a significant correlation of gene signatures with T cell infiltration across a variety of human cancers (Figure 4I). Consistent with our preclinical results in melanoma (B16) and lung cancer (LLC), we observed significant correlation with T cell infiltration in skin cutaneous melanoma (SKCM), lung squamous cell carcinoma (LUSC), and lung adenocarcinoma (LUAD). Genes altered in Yap-cKO TILs were also significantly associated with patient survival across several cancers (Figure 4J), as seen most significantly in LUAD for both CD4+ and CD8+ Yap-cKO gene signatures (Figures 4K-L). These analyses showed that increased Yap activity in TILs correlated with poorer prognosis and lower T cell infiltration, suggesting that Yap function in T cells contributes to aggressive human cancer development and cancer immunosuppression.

## DISCUSSION

We present evidence for an immune inhibitory role for Yap in CD4^+^ and CD8^+^ T cells, and offer the first data for Yap as a negative regulator of T cell tumor infiltration. We found that disrupting Yap activity leads to enhanced T cell activation, augmented potential for differentiation, and increased tumor infiltration. Two major signals are necessary for T cell activation: signaling from engagement of the T cell receptor (TCR) with its cognate antigen:major histocompatibility (MHC) complex, and a second signal from the co-stimulatory receptor CD28 binding to ligands CD80 and CD86 [53, 54]. Through recruitment of key effector kinases, phosphatases, and scaffold proteins, transcriptional programs are induced leading to production of cytokines and cytokine receptors, which importantly include IL-2 and CD25, the α chain of the heterotrimeric high-affinity IL-2 receptor. IL-2 binding to the IL-2 receptor, together with TCR signaling and co-stimulation elicit transcriptional changes resulting in proliferation and differentiation [55]. RNA-Sequencing analysis of WT vs Yap-cKO TILs revealed that CD3, CD28, CD80 and CD86 receptor molecules, kinases, phosphatases, and scaffold proteins are all upregulated with Yap deletion, offering a mechanism for enhanced activation of CD4^+^ and CD8^+^ T cells. These mechanisms are significantly broader than the TGFβ-specific mechanisms previously reported for Yap in Tregs [37], implicating Yap as a broad regulator of T cell responses.

We demonstrated that enhanced CD4^+^ and CD8^+^ T cell activation could be observed *in vitro* using either Yap-cKO cells or treatment with verteporfin, a reported inhibitor of Yap-TEAD activity [56]. Interestingly, the effect of Yap deletion or verteporfin-mediated Yap inhibition induced *in vitro* activation of T cells, but did not significantly impact T cell proliferation. These observations are consistent with observations of increased early activation markers in Yap inhibited cells.

Previous studies have demonstrated proliferation has a sharp, switch-like threshold for T cell receptor signal to elicit proliferation [57], and our experimental conditions provide optimal signaling above this threshold. Expression levels of activation markers have been directly linked to TCR signal strength, such as CD69 levels being directly regulated by affinity and dose of TCR ligand [58] and CD71 levels being directly linked to mTOR activation level [59]. Our observations therefore support a novel conceptual advance: that Yap may link TCR signal strength to negative feedback.

The rapid induction of Yap post-activation suggests that there are no Yap-specific inducers in the tumor, but rather that the signals driving Yap expression are shared canonical signals of T cell activation. These observations are consistent with those made previously by Thaventhiran et al. [27], but contrast a Treg specific role proposed by Ni et al [37]. Our data suggest that Yap promotes a normal negative feedback mechanism during T cell activation similar to inhibitory checkpoint molecules, and inhibition of Yap must be timed before or during T cell activation. Our observations interestingly also indicate that Yap plays a prominent role only following activating signals from the microenvironment. This is highlighted by the large number of genes impacted in Yap-deleted CD4^+^ and CD8^+^ T cells isolated from TILs as compared to TDLNs, and is consistent with our data showing Yap levels increase after T cell activation. These data suggest that therapeutic inhibition of Yap in T cells may have fewer side effects compared to strategies such as checkpoint blockade, since the regulatory activity of Yap is synchronized with T cell activation and appears specific to sites of active T cell priming. Yap connects a variety of extracellular stimuli into intracellular cues that informs the cell of its own structural features (actin cytoskeleton, polarity, cell shape) as well as its location and surroundings (mechanical, adhesion, ECM), instructing cellular survival, proliferation, differentiation and fate. Therefore, a better understanding of how Yap is regulated by these signals in T cells may reveal new insights into immunomodulatory mechanisms.

Regulation of T cell activation by the microenvironment through polarizing cytokines allows for diverse, context-specific differentiation and functional diversity in CD4^+^ T cells [3]. We observe that Yap-cKO CD4^+^ T cells have increased capacity to differentiate towards Th1, Th2, and Th17 phenotypes, implicating Yap in roles beyond Treg regulation. Our observations indicate that Yap does not preferentially bias control of T cell differentiation, but instead permits responses to local microenvironmental cues. This strongly suggests that Yap inhibition may better enable responses to current immunotherapy strategies. Clues into Yap-regulated events are embedded in our RNA-Sequencing of Yap-cKO TILs, which show broad changes in T cell responses. Genes regulated by Yap include several key effectors of T cell activation, including those controlling JAK-STAT signaling that modulate the polarization of naïve CD4s into T helper subsets [60]. Notably, Yap-regulated genes are also enriched for those regulated by the TGF*β* and Wnt pathways, which have important pleotropic roles in T cell biology [61, 62]. Given the known convergence of Yap with these immunomodulatory pathways [63] it is likely Yap directs their transcriptional targets and signal strength. Our study provides novel evidence of CD4^+^ and CD8^+^ T cell-intrinsic effects of Yap on T cell activation and tumor infiltration.

An important finding of our studies is that Yap-cKO mice show delayed tumor growth in conjunction with increased infiltration and expression of effector molecules. The well-characterized B16F10 melanoma and LLC lung cancer models generate tumors that are characterized as immune deserts [41-44, 64-66]. In immune deserts, T cells are completely excluded from the tumor microenvironment due to suppressed tumor immunogenicity and insufficient T cell priming, co-stimulation, and activation [24]. Patients with immune desert tumors show reduced survival, and are not responsive to immune checkpoint inhibitor (ICI) strategies. The ability for Yap-cKO CD4^+^ and CD8^+^ T cells to become activated and infiltrate tumors that normally inhibit T cell infiltration is therefore very notable and important, as these phenotypes are key for combating tumor growth in human cancer patients [67]. Adoptive transfer of Yap-cKO CD8^+^ T cells showed that CD8^+^ T cells have intrinsically enhanced tumor infiltration capacity compared to WT host CD8^+^ T cells into B16F10 tumors. These data are novel and suggest that one of the early events in Yap-mediated tumor immunosuppression may be exclusion of CD8^+^ T cells from the tumor. In mice with Yap-cKO T cells, tumor cell killing by early infiltration of CD8^+^ T cells may elicit 1) tumor antigen release for enhanced intratumoral T cell priming and 2) increased intratumoral activation of CD8^+^ T cells, causing chemokine secretion and recruitment of CD4^+^ T cells and other immune cells. These data provide a new conceptual framework for the sequence of immunosuppressive events orchestrated by Yap in tumors, and indicate that the regulatory effects of Yap are tumor-specific and not systemic. The significant correlation of CD8^+^ and CD4^+^ Yap-cKO gene signatures with tumor T cell infiltration in TCGA data suggest that Yap represses CD8^+^ and CD4^+^ T cell migration and tumor infiltration in human cancers. In particular, our preclinical results in melanoma and lung cancer models were mirrored in human SKCM, LUAD, and LUSC. Low activity of Yap-regulated CD4^+^ and CD8^+^ gene expression signatures correlated with immune infiltration in 19 TCGA cancer types, and strongly correlated with survival in 11 TCGA cancer types, highlighting Yap activity in T cells has broad potential as a target for cancer therapy.

Collectively, our data show that Yap is an important immunosuppressor and suggest that inhibition of Yap activity in T cells could have important clinical implications in T cell therapies against cancer and other diseases. We integrate findings from prior studies and extend them, showing for the first time that Yap is expressed in activated CD4^+^ and CD8^+^ T cells and plays a regulatory role in T cell activation for both subsets. Insights from our study motivate future investigation of Yap inhibition to enhance anti-tumor responses and on the broader role and mechanisms of Yap in regulating T cells in homeostasis and disease.

## MATERIALS & METHODS

### Mouse Strains and Genotyping

Yap-loxP/loxP mice provided by Dr. Jeff Wrana and previously described [68] were backcrossed to the C57BL/6 background for 10 generations, and bred with the Tg(Cd4-cre)1Cwi (Jax: 022071) [69] and LSL-EYFP (Jax: 006148) [70] lines to derive Yap-loxP/loxP; LSL-EYFP; CD4-Cre mice. For adoptive cell transfers, CD4-cre mice were crossed with LSL-tdTomato mice (Jax: 007914) [71]. All experiments were performed with 6-10 weeks old mice. Animal protocols and study designs were approved by Boston University School of Medicine and UMBC. Mice were maintained in pathogen-free facilities at BUMC and UMBC and were PCR genotyped using published protocols [68–71].

### Cell culture and mouse tumor challenges

B16F10 mouse melanoma cells (ATCC CRL-6475) and LLC1 Lewis lung carcinoma cells (ATCC CRL-1642) were cultured in DMEM supplemented with glucose, L-glutamine, sodium pyruvate, 10% FBS, penicillin and streptomycin. Cells were split once they reached 70% confluency and were not used for mouse challenge past a fifth passage. T cells were cultured in RPMI 1640 supplemented with 10% FBS, 1 mM sodium pyruvate, 50 µM β-ME, penicillin, streptomycin, 2 mM L-glutamine, 100 mM non-essential amino acids, 5 mM HEPES free acid and *β*-mercaptoethanol. For tumor inoculations, 5×10^4^ B16F10 cells or 5×10^5^ LLC cells were injected subcutaneously on the right flank of each mouse on day 0. Tumor volume was estimated using the formula (*L* × *W*^2^)/2. Survival endpoint was reached once the tumors measured 500mm^3^, around day 15. Mice were euthanized by isoflurane inhalation and subsequent cervical dislocation and tumors were harvested for further prospecting.

### T cell isolation

Spleens from WT or YAP-cKO mice were pushed through a 70µm mesh (Falcon) using an insulin syringe plunger and washed with PBS. Cells were treated with ACK red blood cell lysis buffer and splenocyte single cell suspensions were prepared for magnetic separation or stained for sorting by flow cytometry. CD4^+^ or CD8^+^ T cell enrichment was performed using magnetic beads (Miltenyi Biotec or STEMCELL Technologies). Naïve CD4^+^ T cells were isolated using a naive CD4^+^ T cell isolation kit (Miltenyi Biotec).

### Adoptive cell transfers

For the adoptive cell transfers, EYFP^+^ Yap-cKO CD8^+^ T cells were mixed 1:1 with dtTomato^+^ WT CD8^+^ T cells and injected intravenously into 8-week-old WT C57BL/6 mice (Taconic) on day 0. Mice also received a subcutaneous injection of 5×10^4^ B16F10 cells on the same day. On day 15 of tumor growth, tumors were harvested and stained with dead cell dye, CD45, CD3, CD4 and CD8 antibodies for flow cytometric analysis.

### Tumor digestion

B16 tumors from WT and YAP-cKO mice were dissected, mechanically disrupted and digested in serum free media containing 2mg/ml collagenase type I (Worthington) and DNase I (Sigma) for 30 minutes at 37°C in a rotator. Tumor digests were then passed through a 70µm mesh (Falcon) using an insulin syringe plunger and washed with PBS. Cells were treated with ACK red blood cell lysis buffer (Gibco) and tumor single cell suspensions were prepared for staining and analysis by flow cytometry. For determining absolute numbers of tumor infiltrating T cells by flow cytometry, AccuCount fluorescent particles (Spherotech) were added to the tumor digests.

### Flow cytometry

Isolated splenocytes or tumor digests were washed with PBS and stained with the LIVE/DEAD fixable near-IR dead cell stain kit (Invitrogen). Cells were then washed with stain buffer (BD), resuspended in stain buffer containing Fc block (BD) and incubated for 5 minutes at 4°C. Surface antibodies were added in predetermined concentrations and cells were incubated for 30 minutes at 4°C or 15 minutes at room temperature, before being washed with BD stain buffer and resuspended in PBS for flow cytometric analysis. For intracellular cytokine staining, cells were fixed and permeabilized using the BD Cytofix/Cytoperm Fixation/Permeabilization kit (BD) after dead cell dye and surface staining. For transcription factor staining, the eBioscience Foxp3/ Transcription factor staining buffer set was used (eBioscience). Flow cytometry analyses were performed on BD LSRII at Boston University School of Medicine Flow Cytometry Facility or at the University of Maryland School of Medicine Center for Innovative Biomedical Resources, Flow Cytometry Shared Service and analyzed by FlowJo (TreeStar).

### T cell activation and proliferation assays

CD4^+^ or CD8^+^ T cells were isolated using magnetic beads as described above. Cells were stained using the CellTrace Violet or CFSE Cell Proliferation Kit (Life Technologies). Briefly, purified cells were washed with PBS and incubated with CellTrace dye for 20 minutes at 37°C protected from light. After 20 minutes, complete RMPI medium was added to the cell suspension and the cells were incubated 5 minutes further before being washed and resuspended in complete RPMI medium. Cells were cultured in 96 well plates at 1×10^5^ cells per well and were stimulated using *α*CD3/CD28 dynabeads (Gibco) at a 1:1 ratio with T cells. On days 1 and 3, cells were stained with dead cell dye as well as lineage and activation markers CD3, CD4, CD8, CD69, CD71, and CD25 and proliferation was measured at the same time.

### CD4^+^ T cell in vitro differentiation

For CD4^+^ T cell in vitro differentiations into Th1, Th2 and Th17 subsets, naïve CD4^+^ T cells were enriched using magnetic beads (Miltenyi Biotec). Purified cells were plated at 1×10^5^ cells per well on a 96 well plate coated with 10μg/ml *α*CD3 (Biolegend), and cultured with 2μg/ml soluble *α*CD28 (Biolegend). The following conditions were specific to each differentiation regime: Th1, 10ng/ml IL-12 and 10μg/ml *α*IL-4; Th2, 50ng/ml IL-4, 10μg/ml *α*IFN*γ* and 10μg/ml *α*IL-12; Th17, 50ng/ml IL-6, 20ng/ml IL-1*β*, 5ng/ml IL-23, 1ng/ml TGF*β*, 12μg/ml *α*IFN*γ* and 10μg/ml *α*IL-4. Cells were cultured for 5 days, before being stimulated with 50ng/ml PMA (Sigma, P1585) and 1μg/ml ionomycin (Sigma, I0634) for 6 hours at 37 °C in the presence of Golgistop (monensin, BD) or Golgiplug (brefeldin, BD) added after the first 30 minutes of stimulation. Cells were stained with dead cell dye, surface markers (CD3, CD4, CD8, CD25) and intracellular cytokines (IFN*γ*, IL-17) or transcription factors (GATA3) as described above.

### Immunofluorescence microscopy

Harvested B16 tumors were fixed overnight in PLP fixative, followed by incubation in 15% and 30% sucrose. Tumors were embedded in OCT and frozen. Cryosections were cut at 5μm thickness and stored at −20°C. On the day of staining, slides were air dried for 1 hour and fixed in acetone for 10 minutes. Slides were washed with PBS and blocked with PBS containing 10% donkey serum, 0.05% sodium azide, 0.5% triton X-100 and 0.2% BSA for 1 hour at room temperature. Slides were stained with rat anti-mouse CD8 (clone CT-CD8a, Fisher) for 2 hours at room temperature, and were subsequently washed with PBS containing 1% Tween. Secondary antibody was diluted in blocking buffer and applied for 1 hour at room temperature (donkey anti-rat alexa 647, Jackson Immuno Research Labs). Slides were washed and mounted in ProLong antifade reagent with DAPI (Life Technologies). Images were acquired using an AxioObserver D1 equipped with a X-Cite 120LED System.

### Quantitative real-time PCR (qPCR)

RNA was extracted using Rneasy Mini Kit (Qiagen), and 1μg was used to generate cDNA using an iScript cDNA Synthesis Kit (Biorad). Taqman primers for mouse GAPDH and YAP (Life Tech) were mixed with cDNA and Taqman Universal Master Mix II (Life Tech) and C_T_ values were normalized to unstimulated controls.

### Sample preparation for RNA-Seq

B16 tumors from WT and YAP-cKO mice were digested as described above. Tumors were subsequently stained with dead cell dye and antibodies against CD45, CD3, CD4, and CD8, and CD4^+^ and CD8^+^ TILs were sorted from each tumor using a BDFACSARIA instrument. CD4^+^ and CD8^+^ T cells were sorted from WT and YAP-cKO tumor draining lymph nodes, as well. Cells were sorted into TRIzol LS reagent (Invitrogen) and RNA was isolated using a miRNeasy micro kit (Qiagen).

### Transcriptomic analyses and gene expression signature extraction

RNA quality was evaluated using Agilent Bioanalyzer 2100 Eukaryote Total RNA Pico chips. RNA-Seq libraries were prepared using the SMART-Seq v4 Ultra Low Input RNA Kit (Takara, 634889) from total RNA, following the manufacturer’s protocol. Libraries were then sequenced on a HiSeq 4000 using 75-bp paired end reads to an average depth of 22,445,650 ± 240,398 reads (SEM). Transcript abundance estimates were quantified using Salmon to mouse reference transcriptome from assembly GRCm38 (mm10), aggregated to gene level for UCSC-annotated genes using tximport, and DESeq2 was used to calculate normalized counts and differential expression [72, 73]. CD4 and CD8 up/down gene signatures were generated through differential expression analysis via DESeq2. Differentially expressed genes (DEG) between CD4 vs. WT and CD8 vs. WT were defined as log2(FC) > 1 (up) or log2(FC) < −1 (down) and FDR < 0.05. The top 25 differentially expressed genes for CD4^+^ and CD8^+^ T cells were visualized with a heatmap combined with a barplot annotation. The heatmap cells represent the log normalized expression values for each sample and include the top 25 upregulated and top 25 downregulated genes. Each row is accompanied by a bar representing the log-fold change in gene expression (KO/WT). Plots were generated using the *ComplexHeatmap* software package available in R.

### Analysis of gene expression signatures in TCGA datasets

The activation of CD4/8 up/down signatures was calculated with Gene Set Variation Analysis (GSVA) [74] in primary tumor samples across multiple TCGA RNA-Seq datasets. Signature activation was summarized by the sum of activation of the upregulated signature and inactivation of the downregulated signature. For example, CD4_sig_ activity = GSVA(CD4_up sig_) – GSVA(CD4_down sig_). Heatmaps for all TCGA analyses were generated with the *pheatmap* software package available in R. TCGA data includes count matrices generated with STAR2/HTSeq downloaded from Genomic Data Commons (GDC) Data Portal. Each count matrix was normalized by relative log expression (RLE) with DESeq2. This analysis was constrained to samples where survival information was available and immune infiltration could be estimated with Tumor Immune Estimation Resource (TIMER) [75]. The average T Cell infiltration per sample was estimated with TIMER in each TCGA dataset, represented as a single heatmap cell. This average was measured separately for CD4 T Cell and CD8 T Cell infiltration. Additionally, these values were summed to observe their additive effect.

For correlation of signatures with T cell infiltration, CD4/8 signature activation was correlated with the sum of CD4^+^ T cell and CD8^+^ T cell infiltration in each TCGA dataset estimated by TIMER [75]. The heatmap is colored by the correlation coefficient and the text of each cell is the adjusted p-value of the correlation. For survival analysis, the CD4/8 signature activation was used to stratify patients across TCGA datasets into high or low activated groups separated by the mean. For each data set, Kaplan-Meier survival plots were generated and the results were summarized with a heatmap. The color of each cell corresponds to the following categories. Dark blue in the graphs signifies the average survival probability for patients with low activity of the signature is higher while red signifies the average survival probability for patients with high activity of the signature is higher. Light blue and light orange signify the same groups with higher survival probability respectively, however the difference is not significant. The text of each cell is the p-value for the survival estimation. The distribution of p-values arising from the multiple survival analyses for each signature across TCGA datasets was compared to a uniform distribution using a Kolmogorov-Smirnov test.

## ACKNOWLEDGEMENTS

We thank Dr. Jeffrey Wrana for sharing the Yap-loxP mice. We acknowledge support from the Boston University Flow Cytometry core and the University of Maryland School of Medicine Center for Innovative Biomedical Resources, Flow Cytometry Shared Service and the Institute for Genome Sciences for RNA sequencing. X.V. was funded by a grant from the NIH National Heart Lung and Blood Institute (R01HL124392). G.L.S. is funded by in part by the UMGCC P30 grant under award number P30 CA134274 from the National Cancer Institute, NIH. EMS was supported in part by the Nathan Schnaper Program (NIH R25CA186872).

## SUPPLEMENTARY FIGURE LEGENDS

**Supplementary Figure 1.**
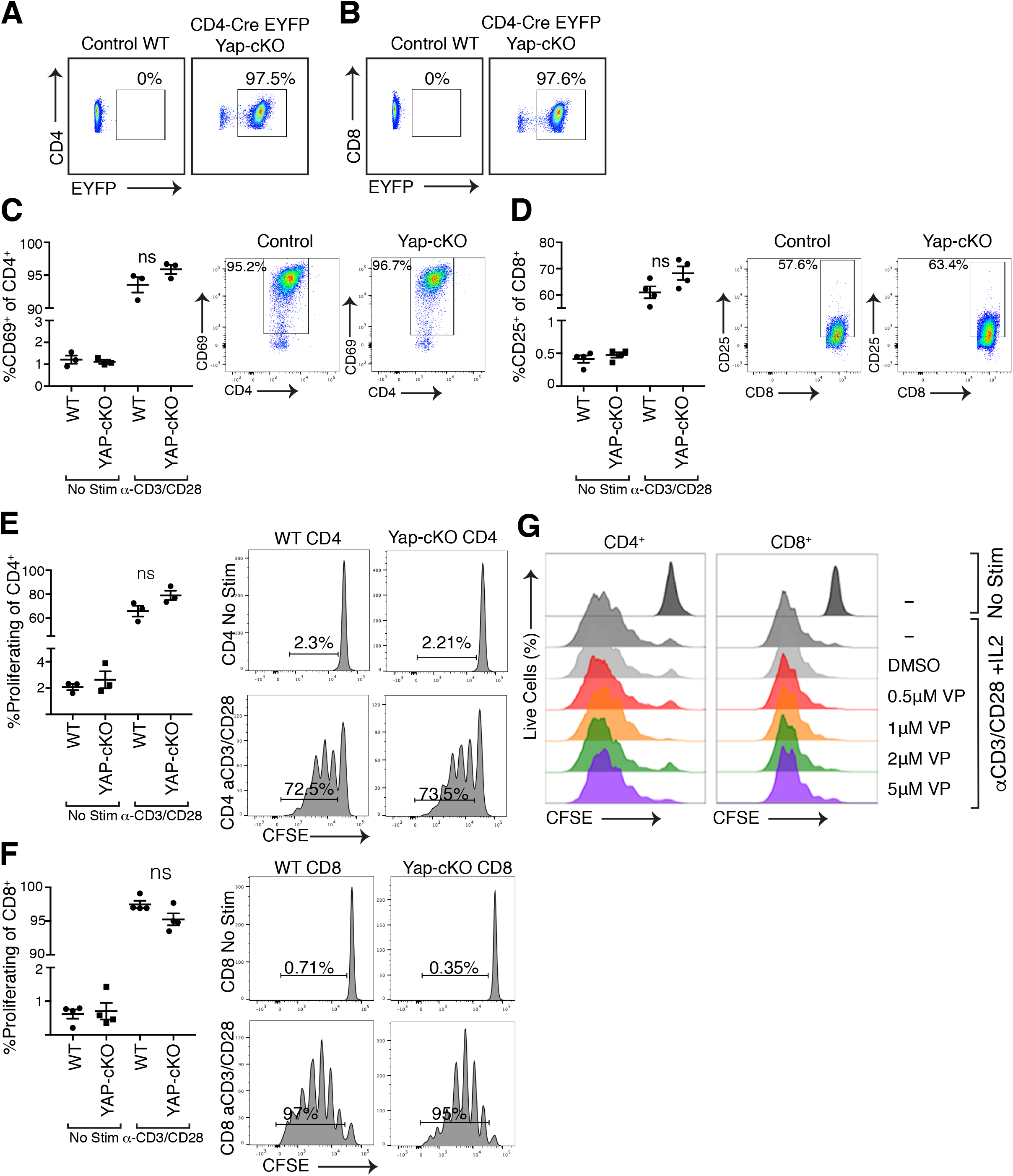
Yap deletion or inhibition does not affect T cell proliferation. CD4^+^ and CD8^+^ T cells were isolated from wild type (WT) or YAP-cKO mice and screened of EYFP expression as well as activation marker expression and proliferation after *α*CD3 and *α*CD28 stimulation. Proliferation was also tested for WT CD4^+^ and CD8^+^ T cells treated with increasing concentrations of Verteporfin under IL-2, *α*CD3 and *α*CD28 stimulation. Statistical differences were determined by using a Student’s t-test, with ns indicating not-significant. a. EYFP expression by flow cytometry on CD4^+^ cells isolated from WT or YAP-cKO mouse spleens. b. EYFP expression by flow cytometry on CD8^+^ cells isolated from WT or YAP-cKO mouse spleens. c. CD69 expression on WT and YAP-cKO CD4^+^ T cells 24 hours post CD3/CD28 stimulation (n=3/group). d. CD25 expression on WT and YAP-cKO CD8^+^ T cells 24 hours post CD3/CD28 stimulation (n=3/group). e. WT vs YAP-cKO CD4^+^ T cell proliferation (n=3/group). f. WT vs YAP-cKO CD8^+^ T cell proliferation (n=3/group). g. Proliferation of DMSO vs verteporfin treated WT CD4^+^ and CD8^+^ T cells (representative of 4 independent experiments).

**Supplementary Figure 2.**
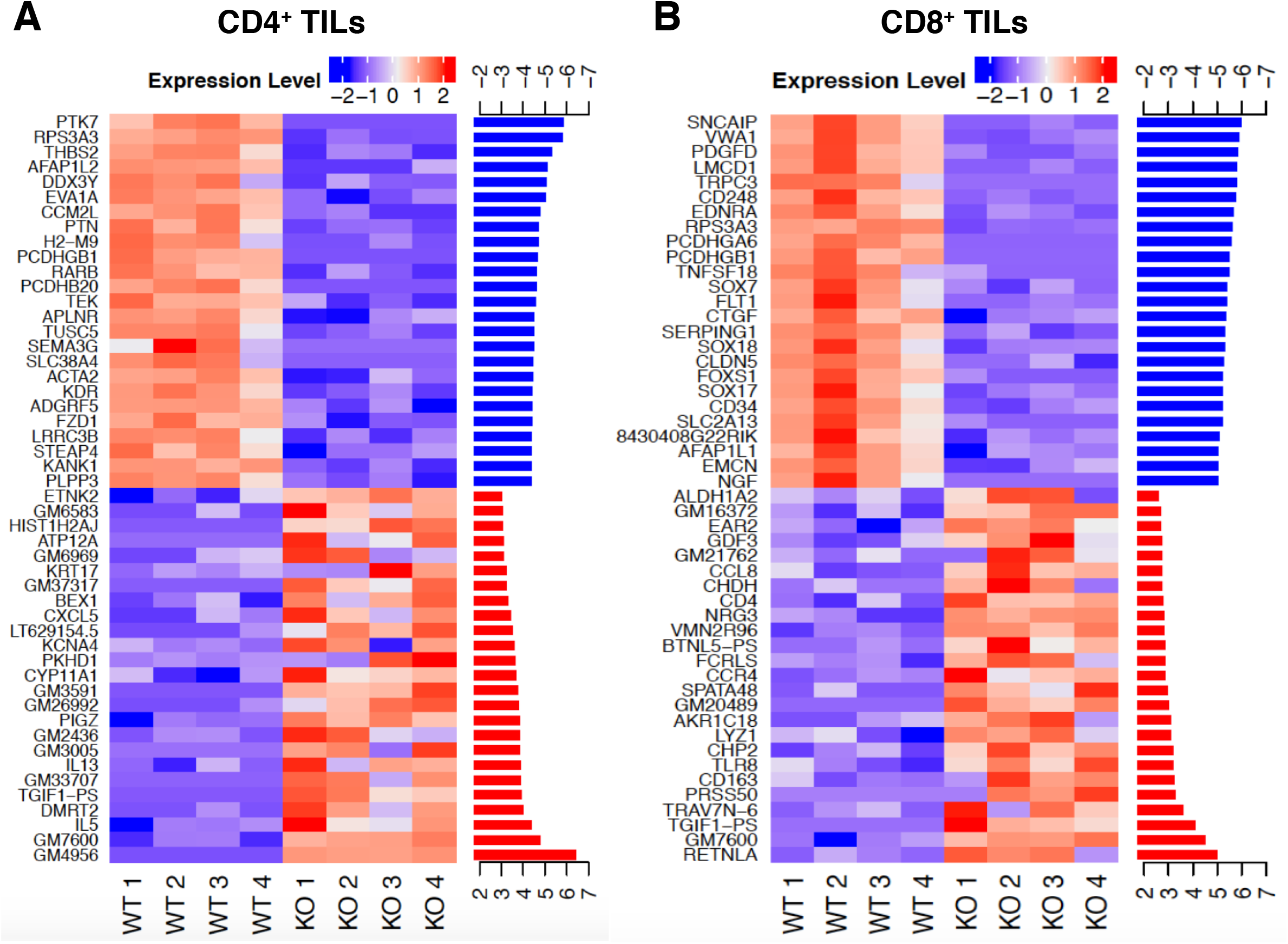
Top 25 upregulated and downregulated genes with YAP deletion in CD4^+^ and CD8^+^ TILs. a. WT vs YAP-cKO CD4^+^ TILs top and bottom DEG heatmap. b. WT vs YAP-cKO CD8^+^ TILs top and bottom DEG heatmap.

**Supplementary Figure 3.**
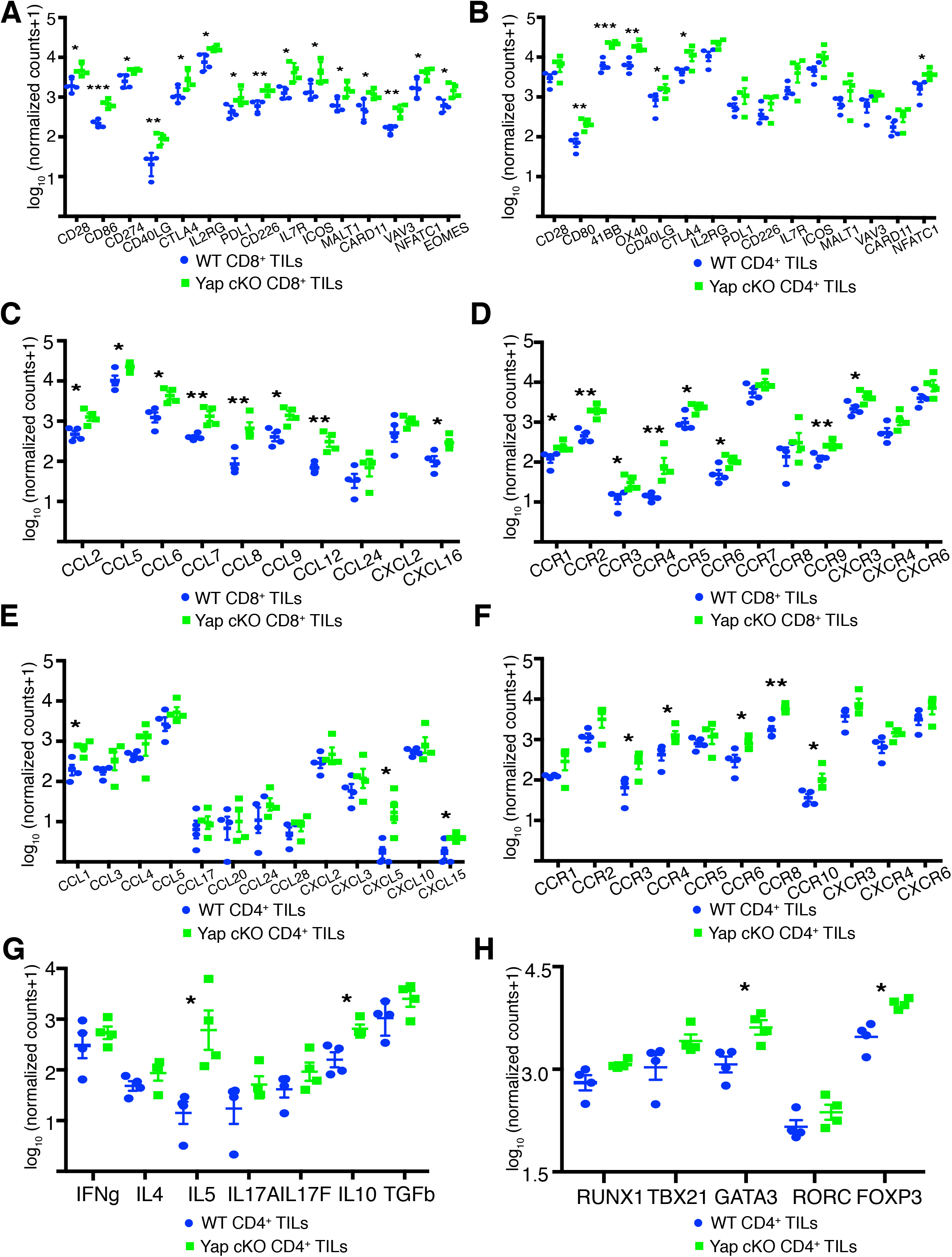
Expression of genes related to T cell activation and co-stimulation, chemokines and chemokine receptors, T helper subset defining cytokines and transcription factors are upregulated in YAP-cKO CD4^+^ and CD8^+^TILs. a. Log_10_(normalized RNA-Seq counts +1) of T cell activation related genes in WT versus YAP-cKO CD8^+^ TILs. b. Log_10_(normalized RNA-Seq counts +1) of T cell activation related genes in WT versus YAP-cKO CD4^+^ TILs. c. Log_10_(normalized RNA-Seq counts +1) of chemokine genes in WT versus YAP-cKO CD8^+^ TILs. d. Log_10_(normalized RNA-Seq counts +1) of chemokine receptor genes in WT versus YAP-cKO CD8^+^ TILs. e. Log_10_(normalized RNA-Seq counts +1) of chemokine genes in WT versus YAP-cKO CD4^+^ TILs. f. Log_10_(normalized RNA-Seq counts +1) of chemokine receptor genes in WT versus YAP-cKO CD4^+^ TILs. g. Log_10_(normalized RNA-Seq counts +1) of T helper subset defining cytokines in WT versus YAP-cKO CD4^+^ TILs. h. Log_10_(normalized RNA-Seq counts +1) of T helper subset defining transcription factors in WT versus YAP-cKO CD4^+^ TILs. Significant differences were determined by a Student’s *t* test; *, p<0.05; **, p<0.01; ***, p<0.001.

**Supplementary Figure 4.**
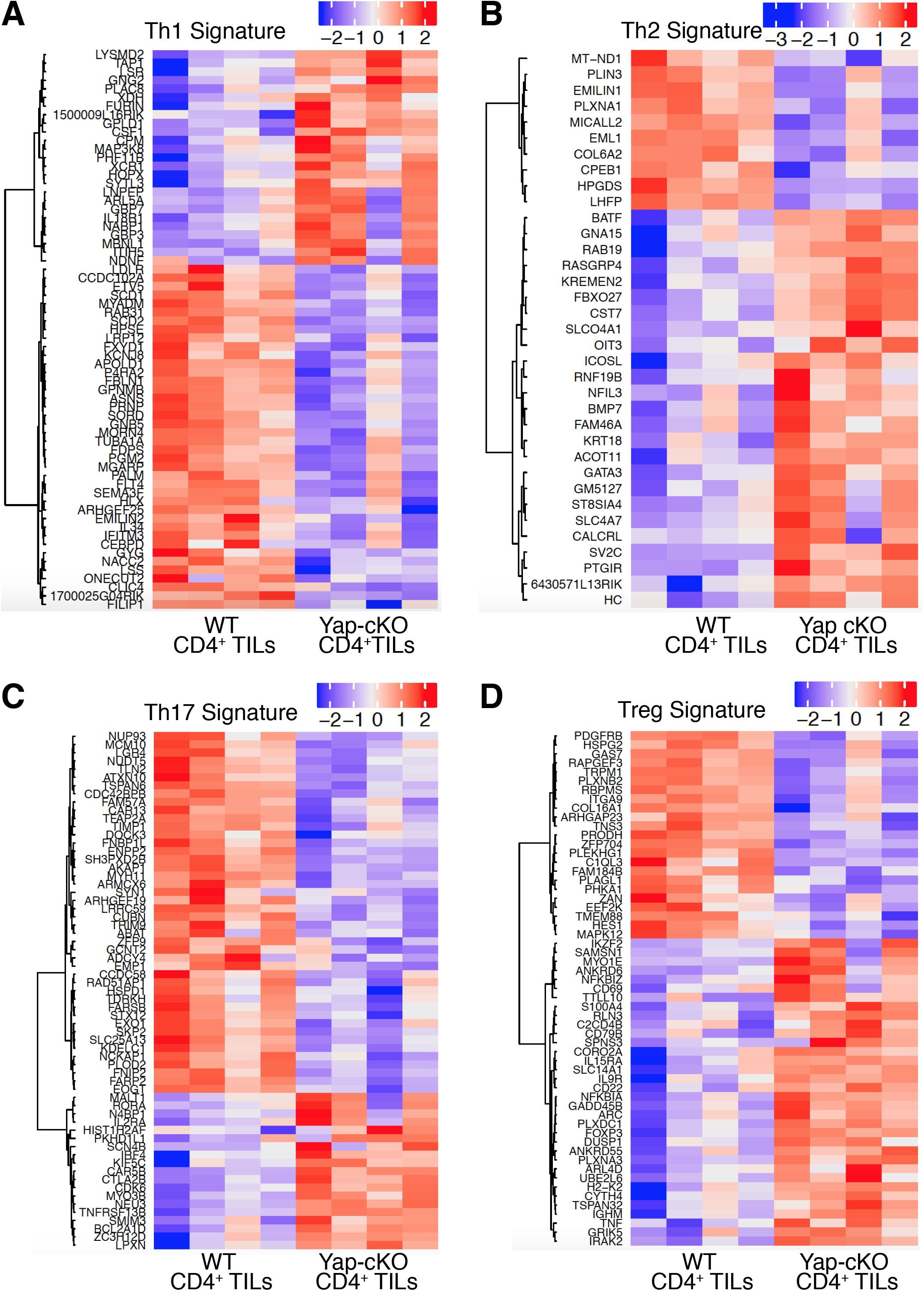
YAP-cKO vs WT CD4^+^ TILs are skewed towards TH2 and Treg signatures compared to WT. a. Heatmap of statistically significant differentially expressed TH1 genes in WT vs YAP-cKO CD4^+^ TILs. b. Heatmap of statistically significant differentially expressed TH2 genes in WT vs YAP-cKO CD4^+^ TILs. c. Heatmap of statistically significant differentially expressed TH17 genes in WT vs YAP-cKO CD4^+^ TILs. d. Heatmap of statistically significant differentially expressed Treg genes in WT vs YAP-cKO CD4^+^ TILs.

